# Transfer Learning for Task Adaptation of Brain Lesion Assessment and Prediction of Brain Abnormalities Progression/Regression using Irregularity Age Map in Brain MRI

**DOI:** 10.1101/345033

**Authors:** Muhammad Febrian Rachmadi, Maria del C. Valdés-Hernández, Taku Komura

**Affiliations:** School of Informatics, University of Edinburgh, Edinburgh, UK; Centre for Clinical Brain Sciences, University of Edinburgh, Edinburgh, UK

**Keywords:** brain lesion’s progression/regression prediction, brain MRI analysis, task adaptation, weakly supervised deep neural networks.

## Abstract

The Irregularity Age Map (IAM) for the unsupervised assessment of brain white matter hyperintensities (WMH) opens several opportunities in machine learning-based brain MRI analysis, including transfer task adaptation learning in the MRI brain lesion’s segmentation and prediction of lesion progression and regression. The lack of need for manual labels is useful for transfer learning. Whereas, the nature of IAM itself can be exploited for predicting lesion progression/regression. In this study, we propose the use of task adaptation transfer learning for WMH segmentation using CNN through weakly-training UNet and UResNet using the output from IAM and the use of IAM for predicting patterns of WMH progression and regression.

## 1 Introduction

Magnetic Resonance Imaging (MRI) facilitates identifying brain pathologies. However, variations in MRI acquisition protocols and scanner manufacturer’s parameters lead to differences in the appearance of the clinical MRI features making their automatic detection challenging. Although the widespread use of MRI has produced large amount of datasets to be used in machine learning approaches, the lack of expert labelled data limits their applicability.

A new method named Irregularity Age Map (IAM) has been recently proposed for detecting irregular textures in T2-FLAIR MRI without requiring manual labels for training [5]. The IAM indicates the degree in which the texture of the neighbourhood around each pixel/voxel differs from the texture of the tissue considered normal. Differently, most machine learning algorithms generate a map indicating the probability of each pixel/voxel of belonging to a particular class (e.g., normal white and grey matter, cerebrospinal fluid, lesions, etc). We believe that the unsupervised nature and the concept of IAM itself are useful for: 1) task adaptation learning in assessing MRI abnormalities and 2) generation of progression/regression patterns that can be used to predict the evolution of these abnormalities. These two topics are the main contributions in this study.

## 2 Task Adaptation Transfer Learning in MRI

### 2.1 Current Approaches of Transfer Learning in MRI

Deep neural networks (DNN) architectures are considered the state-of-art machine learning models in MRI data classification and segmentation as they exhibit or surpass human-level performance on the task and domain they are trained. However, when the domain changes (e.g. imaging protocol or sequence type differ), or they are asked to perform tasks that are related to but not the same task they were trained for (e.g. lesion segmentation vs. lesion assessment), they suffer a significant loss in performance.

Transfer learning (TL) helps dealing with these novel scenarios, as enables a model trained on one task to be re-purposed on a second related task. In DNN, TL is very useful because the first few layers of DNN learn the general visual building of image, such as edges, corners and simple structures while the deeper layers of network learn more complex task-dependent features [1]. Because of that, domain, task or distribution in training and target processes of TL can be different and adjusted to fit the final purpose better.

Domain adaptation TL, where data domains in training and testing processes are different, has been explored in previous studies and shown improving the final performance. In one study, TL has been reported improving Support Vector Machine’s performance in MRI segmentation using different distribution of training data [8]. In other studies, pre-trained DNN using natural images was used for segmentation of neonatal to adult brain images [9] and pre-trained DNN using MRI data from other protocols was used for brain brain lesion segmentation [1].

Whereas, task adaptation TL, where tasks in training and testing processes are different, has not widely explored in medical image analysis. However, the newly proposed unsupervised method of Limited One Time Sampling IAM (LOTSIAM) [5] has been reported to serve the purpose of white matter hyperintensities (WMH) segmentation performing at the level of DNN architectures trained for this specific purpose while executing a different task *i.e.*, extracting irregular brain tissue texture in the form of irregularity age map (IAM).

### 2.2 Weakly-Training CNN in MRI using Age Map

In this study, we explore the use of adapting the task of WMH segmentation on DNN, by using the IAM produced by LOTS-IAM as target instead of binary mask of WMH manually generated by an expert. We evaluate how the DNN recognition capabilities are preserved during the task adaptation TL process.

For our experiments we selected UNet [7] and UResNet [2] architectures used in various natural/medical image segmentation studies. They were also evaluated in the original study of LOTS-IAM [5]. In this study we made two modifications to allow UNet and UResNet to learn IAM: 1) no non-linear activation function (*e.g.*, sigmoid, softmax or ReLU) is used in the last layer of both architectures and 2) mean squared error loss function is used instead of Dice similarity coefficient or binary cross entropy in both architectures.

## 3 Brain Lesion’s Progression and Regression

### 3.1 Prediction of Brain Lesion’s Progression/Regression

Brain lesion’s evolution over time is very important in medical image analysis because it not only helps estimating the pathology’s level of severity but also seelcting the ’best’ treatment for each individual patient [6]. However, predicting brain lesion’s evolution is challenging because it is influenced by various hidden parameters unique to each patient. Hence, brain lesions can appear and disappear at any point in time [6] and the reasons behind it are still unknown.

Previous studies that have modelled brain lesion’s progression/regression, use longitudinal (*i.e*., time-series) data to formulate lesion’s metamorphosis [3,6] by estimating direction and speed of lesion’s evolution over time. Hence, multiple scans are necessary to simulate lesion’s evolution.

In this study, we propose the use of IAM for simulating brain lesion’s evolution (*i.e.*, progression and regression) by using one MRI scan at one time point. This is possible thanks to the nature of IAM which retains original T2-FLAIR MRI’s complex textures while indicating WMH’s irregular textures. Compared to manually produced WMH binary mask by expert or automatically produced probability mask by machine learning algorithm, information contained/retained in IAM is much richer than the others (see Fig. 1).

**Fig. 1.**
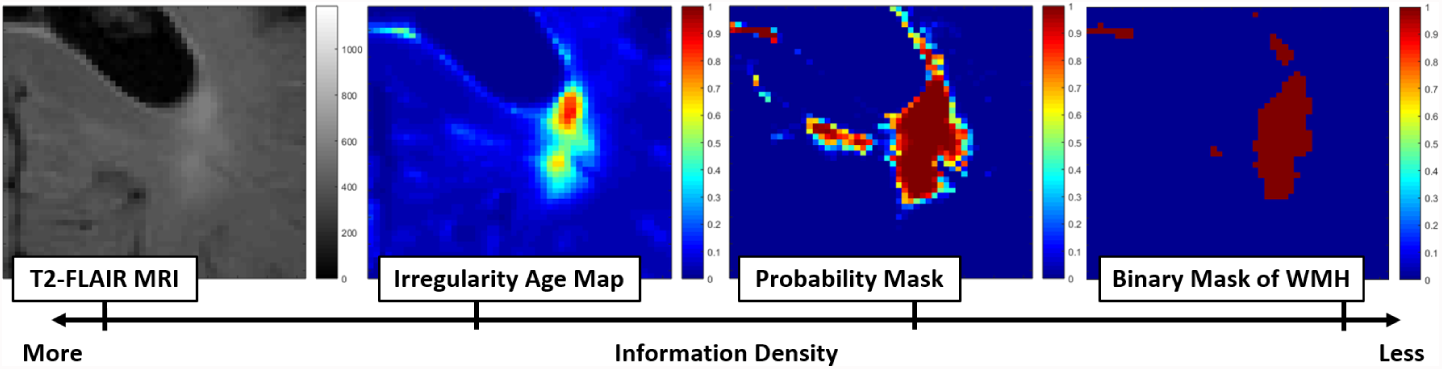
Information density retained in each domain of the original T2-FLAIR, irregularity age map (IAM), probability mask and binary mask of WMH.

### 3.2 Proposed Brain Lesion’s Regression (Shrinkage) Algorithm

We predict the brain lesion’s regression pattern by lowering the threshold value of the IAM. This is possible as each IAM voxel contains different age value. It can be observed in Fig 1 where age values of brain lesion decrease gradually from the border to the centre of each brain lesion. This is not possible using probability masks produced by most machine learning algorithms or binary masks of WMH produced manually by expert where most lesion voxels have flat value of 1.

The algorithm for predicting brain lesion’s regression is detailed in Algorithm 1. To predict the brain lesion regression pattern, we generate pseudo-healthy T2-FLAIR MRI from computing the age map, detailed in Algorithm 2. Nearest neighbour patches of the original IAM patches are decided using the distance value calculated using the distance function as per Equation 1 below.

#### Algorithm 1 Brain lesions regression (shrinkage) prediction algorithm

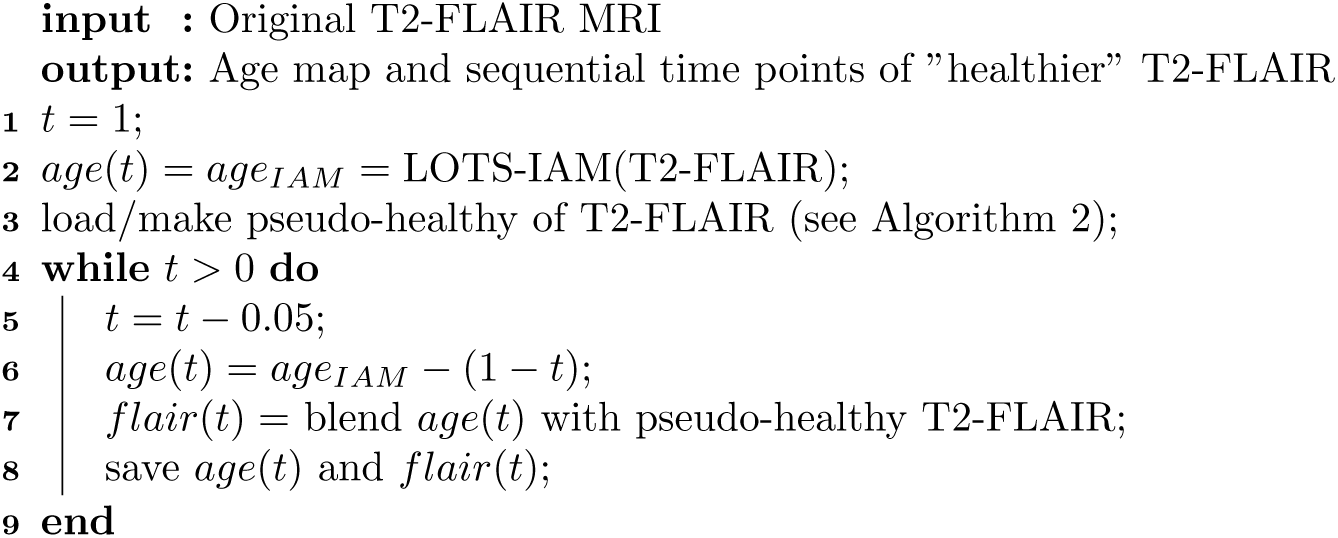

#### Algorithm 2 Pseudo-healthy MRI generation algorithm

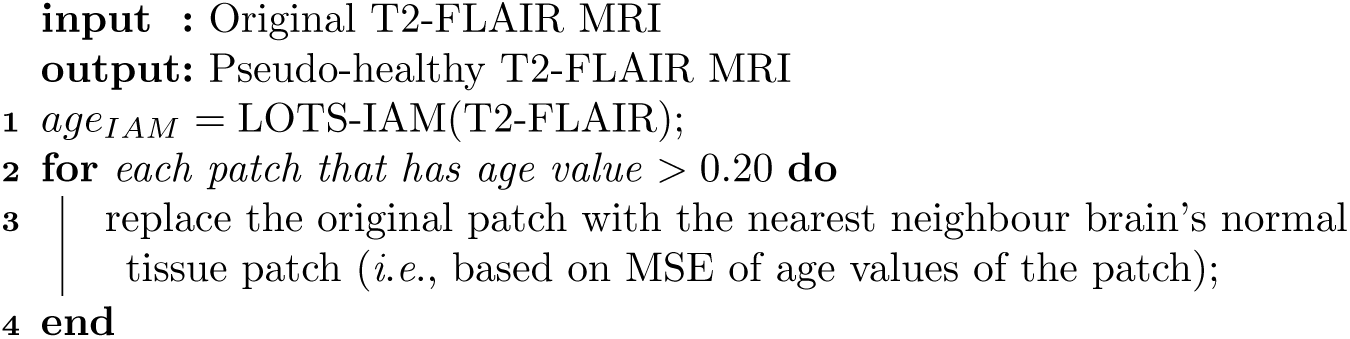

### 3.3 Proposed Brain Lesion’s Progression (Growth) Algorithm

Compared to the previous algorithm for predicting regression, the algorithm for predicting brain lesion progression is more complex as it involves nearest neighbour searching and patch replacement processes. The algorithm still uses IAM produced by the LOTS-IAM. The idea is simple; we need to find similar (*i.e*., nearest neighbour) IAM patches for each original IAM patch but the nearest IAM patch needs to have slightly higher age values than the original IAM patch. Once we find it, the original IAM patch is replaced. Once all patches are replaced by their nearest IAM patches, a new T2-FLAIR MRI showing brain lesion progression can be produced by blending the new IAM with the pseudo-healthy T2-FLAIR MRI.

The algorithm for predicting brain lesion progression is detailed in Algorithm 3. It uses the pseudo-healthy T2-FLAIR MRI produced by Algorithm 2. The distance function used in algorithms 2 and 3 is defined below. Let **s** be the original IAM patch and **t** be the candidate of nearest neighbour patch, the distance *d* between the two patches is:
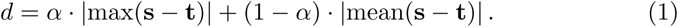

where *α* = 0.5. Whereas, the patch’s size used in this study is 4 × 4.

#### Algorithm 3 Brain lesions progression (growth) prediction algorithm

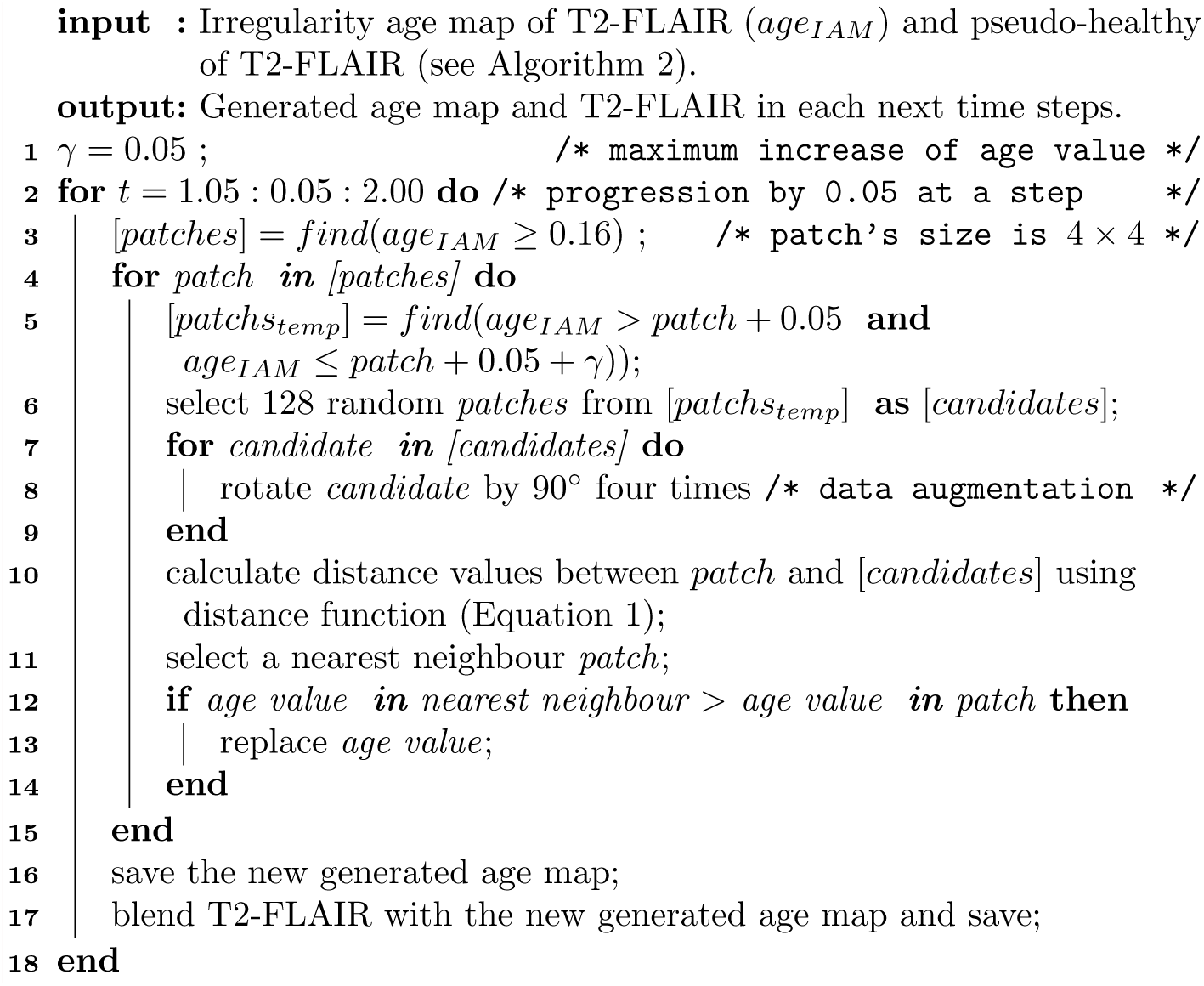

## 4 MRI Data and Experiment Setup

A set of 60 T2-Fluid Attenuation Inversion Recovery (T2-FLAIR) MRI data from 20 subjects was used. The Dice similarity coefficient (DSC) was used to evaluate performance of UNet and UResNet segmenting WMH weakly-trained using IAM. Each T2-FLAIR MRI volume has dimension of 256 × 256 × 35. Data used in this study were obtained from the ADNI [4] public database^1^.

## 5 Results

### 5.1 Weakly-Mraining of UNet and UResNet using IAM

Fig. 2 shows the performance of the two algorithms evaluated in this study: UNet(1) and UResNet(2) segmenting WMH in our sample. Fig. 2A shows the DSC of the WMH masks obtained for the 60 MRI data by both algorithms without TL(Aa) and with TL(Ab) using the results of LOTS-IAM thresholded at 0.18 (see [5]). Both DNN schemes could yield better results if task-adaptation TL using IAM is used. However, the IAM’s dependence on pre-processing poses a risk for their use in TL, as it can also worsen DNN’s performance.

**Fig. 2.**
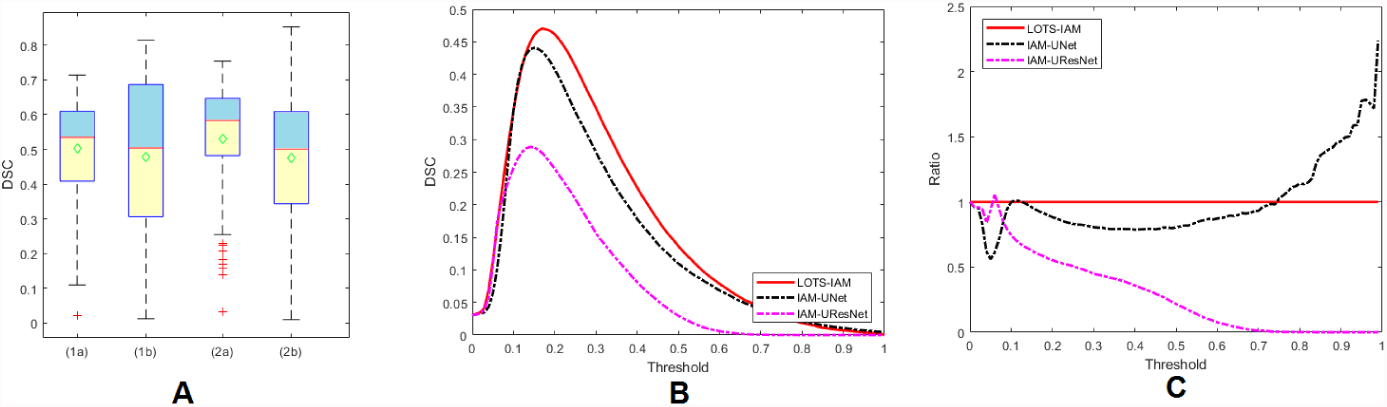
Performance of UNet(A1) and UResNet(A2) in WMH segmentation without transfer learning(Aa) and using transfer learning(Ab, B and C).

As Fig. 2B shows, peak mean performances are 0.4704 (0.1587) for IAM, 0.2888 (0.0990) for IAM-UResNet and 0.4409 (0.1410) for IAM-UNet. The latter performs 15.21% better than the former, which is quite opposite to when TL is not used (see [5]). Our guess is that residual blocks in UResNet perform poorly if it has to learn from real values of IAM. On the other hand, UNet learned from IAM with minimal performance drop (*i.e*., 2.95% drops from the LOTS-IAM) compared with when learned from manual WMH labels (*i.e.*, 6.21% differences as per [5]). Although the performance of IAM-UResNet and IAM-UNet apparently follow the LOTS-IAM’s performance at different thresholds in terms of DSC, a closer look at the learning process shows these relationships are not linear. Fig. 2C shows the ratio between the mean DSC values of these DNN schemes and LOTS-IAM output. The peak DSC performance is not achieved using exactly the same threshold.

### 5.2 Result on Brain Lesion’s Progression/Regression Prediction

Fig. 3 shows visualisations of IAM and T2-FLAIR generated from the original IAM and T2-FLAIR (centre with *t* = 1.00) by using Algorithms 1, 2 and 3. The generated regression step of IAM and T2-FLAIR (2*^nd^* column with *t* = 0.50) was generated by using Algorithm 1 whereas the generated progression steps of IAM and T2-FLAIR (4*^th^* and 5*^th^* column with t = 1.25 and *t* = 1.50) were generated by using Algorithm 3. On the other hand, the pseudo-healthy T2-FLAIR (1^st^ column with *t* = 0.00) was generated by using Algorithm 2.

**Fig. 3.**
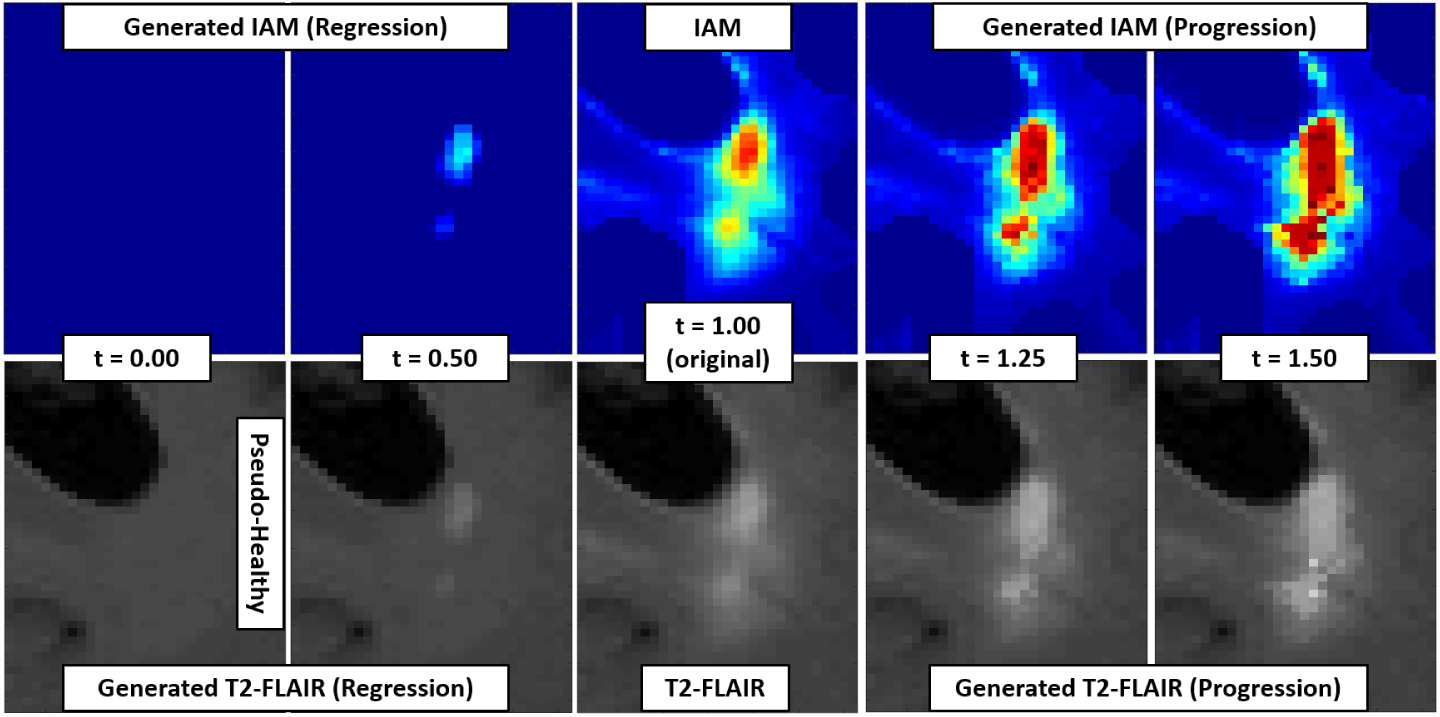
Visualisation of brain lesion’s progression and regression prediction by manipulating age values of IAM.

As Fig. 3 shows, prediction of brain lesion’s regression works really well for WMH, but prediction of brain lesion’s progression shows a small unmatched tessellation problem. This problem is common in computer graphics when an old patch needs to be replaced with a new one, so it should be easy to fix in the next iteration of this experiment. Nevertheless, this experiment shows the suitability of IAM for predicting brain lesion’s progression/regression.

## 6 Discussion

In this study, we have demonstrated the use of IAM produced by the LOTS-IAM for weakly-training two DNN schemes, *i.e.*, UResNet and UNet, and predicting brain lesion’s progression/regression. Performance of UNet weakly-trained using IAM is close to the LOTS-IAM and UNet trained by using manual label of WMH and can sometimes be improved. In the future, we will widen our sample and investigate the conditions under which TL improves/worsens the quality of the DNN outputs.

Furthermore, IAM has been shown in this study to be very useful for the brain lesion’s progression/regression prediction. There are still some problems in the prediction of progression such as unmatched tessellation and changing of contrast of the T2-FLAIR. However, it does not change the fact that the use of IAM facilitates the prediction of brain lesion’s progression/regression. Next steps in this research topic would be fixing unmatched tessellation avoiding the effect caused by contrast differences, and the use of pre-trained DNN (*e.g.*, UNet) for predicting brain lesion’s progression/regression.

## Acknowledgement

Funds from Indonesia Endowment Fund for Education (LPDP) of Ministry of Finance, Republic of Indonesia and Row Fogo Charitable Trust (Grant No. BRO-D.FID3668413) (MCVH) are gratefully acknowledged. Data collection and sharing for this project was funded by the Alzheimer’s Disease Neuroimaging Initiative (ADNI) (National Institutes of Health Grant U01 AG024904) and DOD ADNI (Department of Defense W81XWH-12-2-0012).

## Footnotes

1 Database can be accessed at http://adni.loni.usc.edu. A complete listing of ADNI investigators can be found at http://adni.loni.usc.edu/wp-content/uploads/howtoapply/ADNIAcknowledgementList.pdf

## References

1. Ghafoorian, M., Mehrtash, A., Kapur, T., Karssemeijer, N., Marchiori, E., Pesteie, M., Guttmann, C.R., de Leeuw, F.E., Tempany, C.M., van Ginneken, B., et al.: Transfer learning for domain adaptation in mri: Application in brain lesion segmentation. In: International Conference on Medical Image Computing and Computer-Assisted Intervention. pp. 516–524. Springer (2017)

2. Guerrero, R., Qin, C., Oktay, O., Bowles, C., Chen, L., Joules, R., Wolz, R., Valdés-Hernández, M., Dickie, D., Wardlaw, J., et al.: White matter hyperintensity and stroke lesion segmentation and differentiation using convolutional neural networks. NeuroImage: Clinical 17, 918–934 (2018)

3. Hong, Y., Joshi, S., Sanchez, M., Styner, M., Niethammer, M.: Metamorphic geodesic regression. In: International Conference on Medical Image Computing and Computer-Assisted Intervention. pp. 197–205. Springer (2012)

4. Mueller, S.G., Weiner, M.W., Thal, L.J., Petersen, R.C., Jack, C., Jagust, W., Trojanowski, J.Q., Toga, A.W., Beckett, L.: The alzheimer’s disease neuroimaging initiative. Neuroimaging Clinics of North America 15(4), 869–877 (2005)

5. Rachmadi, M.F., Hernández, M.V., Li, H., Guerrero, R., Zhang, J., Rueckert, D., Komura, T.: Limited one-time sampling irregularity age map (lots-iam): Automatic unsupervised detection of brain white matter abnormalities in structural magnetic resonance images. bioRxiv p. 334292 (2018)

6. Rekik, I., Allassonnière, S., Carpenter, T.K., Wardlaw, J.M.: Using longitudinal metamorphosis to examine ischemic stroke lesion dynamics on perfusion-weighted images and in relation to final outcome on t2-w images. NeuroImage: Clinical 5, 332–340 (2014)

7. Ronneberger, O., Fischer, P., Brox, T.: U-net: Convolutional networks for biomedical image segmentation. In: International Conference on Medical image computing and computer-assisted intervention. pp. 234–241. Springer (2015)

8. Van Opbroek, A., Ikram, M.A., Vernooij, M.W., De Bruijne, M.: Transfer learning improves supervised image segmentation across imaging protocols. IEEE transactions on medical imaging 34(5), 1018–1030 (2015)

9. Xu, Y., Géraud, T., Bloch, I.: From neonatal to adult brain mr image segmentation in a few seconds using 3d-like fully convolutional network and transfer learning. In: Image Processing (ICIP), 2017 IEEE International Conference on. pp. 4417–4421. IEEE (2017)

